# Using Time Dependent Rate Analysis to Evaluate the Quality of Machine Learned Reaction Coordinates for Biasing and Computing Kinetics

**DOI:** 10.1101/2025.07.03.662600

**Authors:** Nicodemo Mazzaferro, Suemin Lee, Pilar Cossio, Pratyush Tiwary, Glen M Hocky

## Abstract

Having an accurate reaction coordinate (RC) is essential for reliable kinetic characterization of molecular processes, but there are few quantitative metrics to evaluate RC quality. In this study, we consider the dimensionless *γ* metric from the Exponential Average Time-dependent Rate (EATR) method, which represents the fraction of a biasing potential along the RC that contributes to increasing the rate constant. We demonstrate that *γ* can be used to test whether the utility of a reaction coordinate for predicting kinetics with a Meta-dynamics bias improves as the coordinate is iteratively updated to include new data. We evaluate reaction coordinates approximated via the iterative State Predictive Information Bottleneck (SPIB) approach, which was previously shown to be accurate across six protein–ligand dissociation systems. For these same systems, we compute *γ* values and mean accelerated times 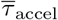. After systematically scanning over fitting parameters, the results show that *γ* increases closer to 1, while 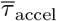 decreases, revealing a consistent inverse correlation. These results demonstrate that *γ* serves as a practical criterion for RC evaluation and offers guidance for selecting SPIB-derived coordinates yielding quantitative kinetic predictions.

## I. INTRODUCTION

Determining biomolecular kinetics is crucial in many areas of biochemistry and biophysics, such as protein folding, ligand unbinding, membrane permeation, and other cellular events. Molecular dynamics (MD) simulations offer the promise of being able to predict kinetics of such processes from simple physical principles.^1–3^ In practice, however, the dynamical processes of interest are “rare events,” the typical timescales of which are dominated by crossing over high free energy barriers, and also dictated by the need to diffuse throughout a metastable state before escaping. Consequently, direct MD simulations that access the timescales of microseconds are rarely sufficient to sample enough configurational space to estimate the transition times of key biological processes, which can range from microseconds to seconds. ^4,5^

To address the sampling problem, a wide range of enhanced-sampling approaches based on adding additional bias to a set of collective variables (CVs), most notably umbrella sampling, metadynamics (MetaD), and on-the-fly probability enhanced sampling (OPES) have been developed to accelerate exploration.^6–9^ After sampling with an applied bias along one or several CVs, equilibrium averages in the unbiased ensemble can be obtained by reweighting.

A separate challenge arises when one is interested in computing dynamical quantities dominated by rare events. Because enhanced-sampling approaches accelerate exploration, they also distort the underlying kinetics of the simulated system. Surprisingly, it is still possible to extract meaningful kinetics from CV-based biasing methods with several approximations. Approaches for doing so include hyperdynamics, infrequent meta-dynamics (iMetaD), OPES flooding, and applying the Kawasaki relation.^10–13^ While many of these approaches have been carefully validated against model and real world systems,^14^ most of these methods require a good CV for biasing that approximates the true reaction coordinate (RC). This is because applying a bias potential along a poor CV has a less understood effect on the slow transition of interest, which must be accounted for. Thus identifying good CVs remains a critical bottleneck, especially for complex systems with many degrees of freedom.

A good biasing variable that approximates the RC should both distinguish the metastable basins of interest and characterize the dynamics of the slow collective modes of a molecular process of interest. The most ideal RC is considered to be the ‘committor’, a function that reports the probability of ending within a product or reactant state at some later time for a current configuration. It is thus the best variable to describe a transition process, as it separates the metastable states of interest by definition and includes the kinetic information necessary to estimate rates.^15–18^ In practice, despite recent progress to compute the committor using machine learning approaches^19–22^, the committor is still difficult to obtain, and it is not easily generalizable across different conditions.^15^ Instead of using the committor as a direct RC, many other approaches have been proposed to learn an approximate RC, including transition-path sampling, diffusion maps, Linear Discriminant Analysis, the variational approach to conformational dynamics, time-lagged independent component analysis, VAMPnets, SGOOP, and others.^23–32^ Building upon these methods, recent studies have shown that incorporating deep learning approaches can improve on RC discovery.

In particular, dimensionality-reduction approaches integrated with deep neural networks have shown much success.^31–34^ However, compressing a high-dimensional process into a single coordinate can lead to the loss of information to accurately describe the reaction mechanism. This loss of information is a major obstacle in extracting a good RC and building a reliable kinetic model. Tiwary and coworkers addressed this challenge by performing an iterative process through the State Predictive Information Bottleneck (SPIB) framework,^35^ to learn the RC associated with kinetic rates.^36^ SPIB employs a deep neural network that identifies a low-dimensional representation of the system’s high-dimensional dynamics that is maximally predictive of the future state of the system after a specified lag time. This approach ensures that the learned RC captures the most predictive features of the system’s slow dynamics, effectively approximating aspects of the committor function. By retraining the network after each round of accelerated sampling, it is believed that SPIB progressively refines the RC. In other words, this proposed algorithm, using SPIB, learns from newly sampled trajectories and then guides subsequent simulation rounds toward more relevant features. Here, we ask the following question: Does this iterative cycle genuinely improve the RC?

For rare events such as ligand dissociation, it is difficult to determine the quality of a collective variable since we do not typically have the true RC to compare with. These considerations raise a broader issue: How can we systematically evaluate the quality of RCs for obtaining rates from biased simulations? For unbiased simulations, one way to determine whether an RC captures the rare event of interest is by performing a Kolmogorov-Smirnov test and using the p-value to determine if the transition time distribution is Poissonian.^37^. Another way is to calculate the committor probability as a function of the RC to ensure that the reactant and product states are well-separated.^23^ We can also rank RCs using the Bayesian transition-path probability *p*(TP | *x*), as has been demonstrated by Best and Hummer.^17^ However, applying these metrics requires unbiased simulation data that includes hundreds of transition events, which is often a great luxury to have for a rare event such as ligand dissociation. To evaluate the RC quality in our biased simulations, we instead use the exponential average time-dependent rate (EATR) method,^38^ which provides the dimensionless metric *γ* that was originally introduced in the Kramers-time dependent rate (KTR) approach^39^. This *γ* quantifies how the RC influences transition times and thus captures how well the bias potential is applied along the RC in the rate-determining directions. When *γ* = 1, the bias acts along the true transition path and has the greatest effect on the transition times, indicating a perfect RC; when *γ* = 0, the bias acts orthogonally to the transition path, indicating a poor RC. In this manner, *γ* serves as a quantitative benchmark of RC performance (see Sec. II C for details). This SPIB + EATR pair, therefore, offers a self-contained, inexpensive, and quantitative solution to observe improvement during RC training. In this context, our study is driven by two central questions: *i*) Does the iterative SPIB procedure enhance the RC for kinetic studies performed here using metadynamics, and *ii*) Can the EATR metric *γ* reliably quantify the quality of any candidate RC? Our results answer both questions affirmatively: *γ* offers a sensitive benchmark for RC comparison, and each round of SPIB refinement leads to improvement in RC quality for more accurate kinetic predictions.

The remainder of the paper is organized as follows. In Sec. II, we review the background on SPIB, infrequent metadynamics (iMetaD), and EATR. We then describe the workflow for assessing RC quality using the EATR *γ* values for the SPIB RC iterations. In Sec.III, we report the EATR *γ* values for the iterations of the SPIB RC and present further analysis on the *γ* metric. Finally, in Sec. IV, we summarize our key findings, discuss the limitations, and propose future directions.

## II. BACKGROUND AND METHOD

In this section, we briefly review the concepts of SPIB,^35^ iMetaD,^11^ and the EATR^38^ *γ* parameter. We then present our integrated workflow, detailing how SPIB-derived reaction coordinates guide iMetaD simulations and how EATR is subsequently applied to quantify RC quality (Fig. 1).

**FIG. 1:**
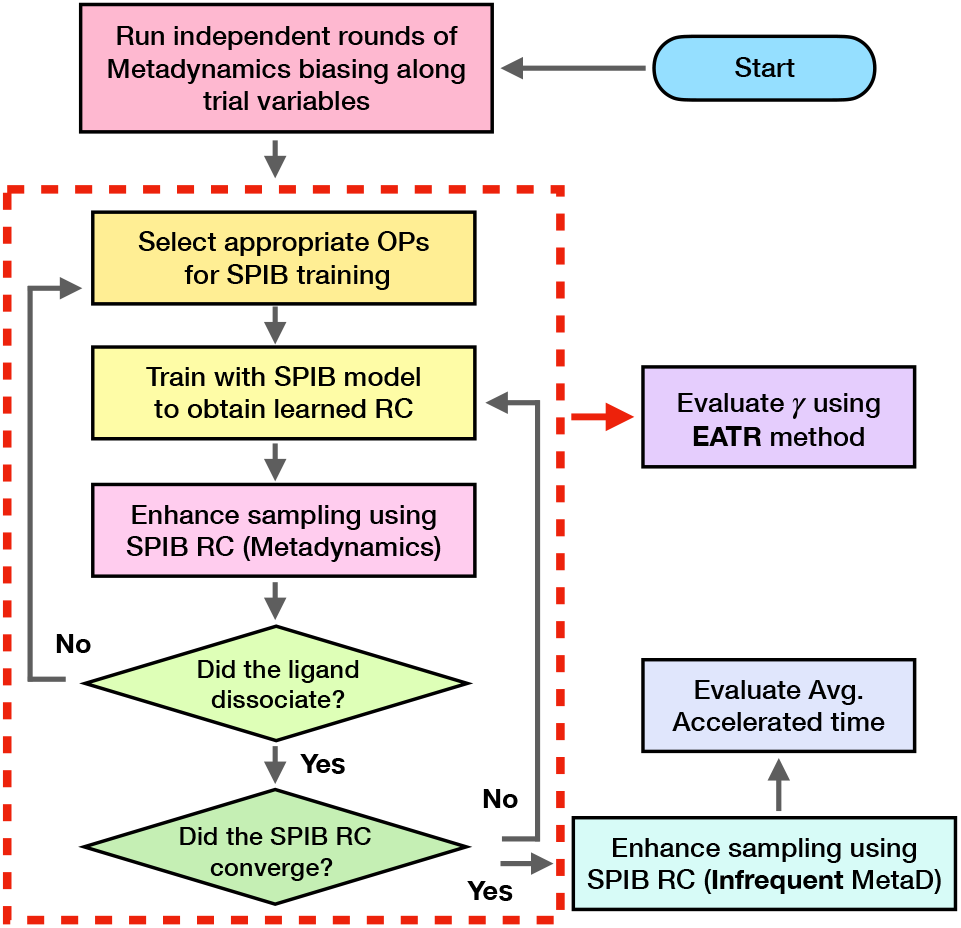
Workflow of SPIB combined with EATR. The gray arrows in the protocol indicate the original SPIB–iMetaD workflow used to compute the residence time of protein–ligand complexes. In Ref. 36, we first performed initial trial runs of WT-MetaD biasing along trial order parameters (OPs). We then iteratively applied SPIB and metadynamics until the SPIB RC of the accelerated time quantitatively reduced. Finally, we ran iMetaD biasing along the converged RC. For each RC at every SPIB iteration, we conducted an additional step of *γ* evaluations, which is indicated as a red arrow. The red dotted box highlights the iterative process of the SPIB.

### A. State Predictive Information Bottleneck (SPIB)

The SPIB method approximates a low-dimensional RC that preserves the metastable dynamics present in the system’s original high-dimensional space.^35^ SPIB is inspired by another information bottleneck-based framework, Reweighted Autoencoded Variational Bayes for Enhanced Sampling (RAVE)^40,41^, but improves upon this formulation. The SPIB approach has demonstrated notable success in learning effective RCs across diverse applications, ranging from crystal nucleation and ligand permeation through lipid membranes to protein conformational changes.^36,42–46^ SPIB CVs can be used with the PLUMED open-source enhanced sampling library^47^, and a tutorial for training and using SPIB CVs is available from the PLUMED-tutorials resource^48^.

SPIB determines a low-dimensional RC that effectively predicts the future metastable states of the system while relying on minimal information from the current configuration. SPIB uses raw MD trajectory data through an encoder-decoder network framework, where the encoder compresses the input into a bottleneck variable that minimizes mutual information with the input, and the decoder maximizes mutual information between the bottleneck and future states after a time lag Δ*t*. By trading off the minimization of input information and the maximization of future predictive information, SPIB simultaneously learns the RC and discovers the location and number of metastable states on-the-fly. A mixture-of-Gaussians prior is used for the bottleneck variable, where the number of Gaussians automatically adjusts to equal the number of metastable states. We emphasize that the number of metastable states in a system is not an intrinsic property of the system, but depends on the time resolution being used to filter out the dynamics. In this spirit, the number of metastable states learned in SPIB decreases as a function of the time-lag parameter.

A notable success of the SPIB framework is its use in protein–ligand dissociation kinetics. In our previous work of Ref. 36, we implemented an iterative SPIB pipeline where each newly learned RC guides a subsequent round of iMetaD (see below Sec. II B), and the resulting trajectories are used to retrain SPIB. This iterative process not only accelerates the observation of the unbinding process but also demonstrates success in SPIB refinements to a more descriptive RC, yielding more accurate dissociation-rate constant estimates.

### B. Infrequent Metadynamics (iMetaD)

iMetaD is a variant of well-tempered metadynamics (WT-MetaD) introduced to evaluate kinetic rates from molecular dynamics (MD) simulations.^7,11^ Like standard MetaD, iMetaD applies a history-dependent Gaussian bias along chosen collective variables (CVs) to accelerate barrier crossings on the free-energy surface.^7,8,49,50^ However, the bias deposition frequency is chosen to be much lower than is typically used to evaluate free energies, such that it avoids depositing the bias on the transition states, and the method preserves the transition state and permits a reliable estimate of kinetics from the biased trajectory. iMetaD relies on the assumption that the selected RC captures the true transition^11^. Under this assumption, iMetaD uses ideas originally pioneered by Voter and Grubmüller^10,51^, and proposes a time-dependent acceleration factor:

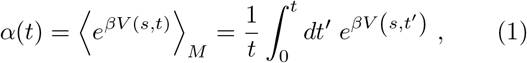

where *V* (*s, t*^′^) is the bias potential at CV value *s* and time *t*^′^, *β* = (*k*_*B*_*T*)^−1^, and ⟨·⟩_*M*_ denotes an average over the metadynamics trajectory of length *t*. Multiplying the physical simulation time *t* by *α*(*t*) yields the accelerated time

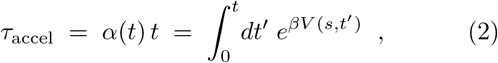

which can be directly compared to experimental or unbiased timescales to recover true kinetic rates. This careful control of bias-deposition frequency is what makes iMetaD uniquely suited for quantitative kinetic studies.

In its original formulation, the mean first passage time for a process (which for a unimolecular process like dissociation is the inverse of the rate constant) is computed as an average over *N* separate iMetaD runs,

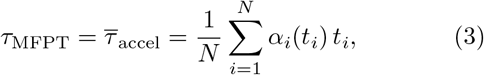

where the bar represents the average over simulations and *t*_*i*_ is the transition time in the *i*th iMetaD simulation. Later, it was shown that it can be more accurate to estimate this time by fitting the empirical cumulative distribution function of rescaled transition times *α*_*i*_*t*_*i*_ to the Poisson model *C*(*t*) = 1 − *e*^−*t/τ*^, where a Kolmogorov–Smirnov test can be applied to assess the validity of the Poisson model.^37^

### C. Exponential Average Time-Dependent Rate (EATR)

The KTR method expanded on the iMetaD method, introducing a more complex form of the cumulative distribution function that also included a second fitting parameter, *γ*.^39^ The KTR method estimates the unbiased transition rate constant *k*_0_ by fitting a time-dependent formulation for the increasing rate constant due to iMetaD biasing, using a relationship based on Kramers’ rate theory

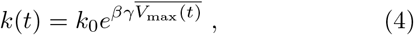

where *k*(*t*) is the biased rate constant at simulation time *t, k*_0_ is the true unbiased rate constant, *V*_max_(*t*) is the maximum value of the bias potential at *t*, and *γ* ∈ [0, 1] is a fitting parameter which scales effect of the bias potential. The overhead bar in Eq. 4 indicates an average over all simulations at time *t. The parameter γ was introduced to account for the use of non-ideal RCs*, with the intuition being that only some of the applied MetaD bias on a CV or set of CVs goes into promoting a transition.

In Ref. 38, it was shown that the usual iMetaD and KTR estimators can be derived through maximum likelihood estimation on biased data from the survival function *S*(*t*), which is the probability for a system to not transition before time *t*, using different forms of a time-dependent rate function *k*(*t*) = *k*_0_*f* (*t*),

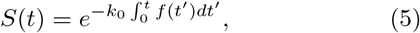

where *f* (*t*) is the “rate scaling function.”

While KTR gives accurate results for real systems, it did not precisely agree with the true unbiased result for a perfect RC.^38^ The EATR method was proposed to correct this. It estimates the transition rate from biased simulations using the relationship,

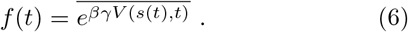

This form is equivalent to iMetaD when *γ* = 1.

In Ref. 38, it was shown that while traditional iMetaD gives a good estimate of transition times for slow biasing, EATR gives a much more accurate estimate for faster biasing for both good and bad RCs *using the same data*. Moreover, *γ* was consistent with the intuition of which RCs would be expected to be good and bad for both a coarse-grained and an all-atom example of protein folding kinetics. Because *γ* should correspond to how effective the bias is in increasing the transition rate, and biasing good RCs affect the transition times more than biasing poor RCs, *γ* is ultimately a measure of the quality of an RC for predicting kinetics.

### D. Overall workflow

Building on the background described above, we now present an integrated SPIB–EATR framework for quantifying RC quality across multiple protein–ligand dissociation systems. We examine six distinct protein-ligand dissociation systems where residence times span approximately 12 orders of magnitude from ∼10 ns to ∼10^3^ s. The systems studied are FKBP–DMSO, FKBP–DSS, T4 Lysozyme–benzene, ABL kinase wild-type (WT) bound to Imatinib, and two ABL kinase mutants (N368S and L364I) bound to Imatinib. Biased trajectories of SPIB iterations for each system were taken directly from Ref. 36, with additional data added for some cases indicated below. All data were generated using GROMACS^52^ and PLUMED^47^ using the protocol described in Ref. 36.

The overall workflow, summarized in Fig. 1, begins with well-tempered metadynamics along trial order parameters (OPs) chosen by chemical intuition. The gray arrow in the workflow diagram denotes the steps originally described in Ref. 36. Trajectories from this initial metadynamics run seed the first SPIB training round; the RC learned in that round is then used to bias along new MetaD simulations. This iterative cycle repeats until the average accelerated time no longer increases. A final set of iMetaD simulations are carried out along the converged RC. Detailed choices of OPs, SPIB hyperparameters, and MD setup were used as reported in Ref. 36.

Next, we apply the EATR analysis to these simulations. For each system, we extract the biased trajectory data from each SPIB iteration, typically eight trajectories per iteration, and compute the mean accelerated time denoted as 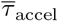. We then use the corresponding bias potentials to evaluate the *γ* metric at each iteration, thereby quantifying RC performance. In the following section, we present these numerical results, describe the EATR parameterization, and discuss how the *γ* values inform RC quality assessment.

## III. RESULTS

In this section, we present two main results. First, we analyze how the RC produced by SPIB at successive iterations relates to the EATR metric *γ*. Second, we provide brief mathematical insights on correlations between the *γ* and the average accelerated time. Finally, we demonstrate that these results are not strongly dependent on the rate at which bias is applied to a particular RC.

### A. Evaluation of SPIB RC via EATR *γ*

To quantify how successive SPIB iterations refine the RC, we first computed the EATR-derived metric *γ* at every iteration for the six protein–ligand systems examined.^36^ In the original EATR approach, both the experimental rate constant *k*_0_ and *γ* are treated as free parameters. Here we compared two fitting protocols. In the first approach, the experimental rate constant *k*_0_ was held fixed at the value reported in Appendix Table T1 for each system. This allows us to fit the survival function only to a single parameter *γ*, which makes the fitting more reliable by removing the possible correlations between *γ* and the rate constant *k*_0_. Although this approach would not apply to all cases, we use this single parameter approach to assess whether SPIB RC quality improves across iterations. We also applied our general EATR approach to fit both *γ* and *k*_0_, with qualitatively similar results for whether the CV is improving, but much lower *γ* values showing there is likely still more that can be done to get more efficient rates as discussed in the conclusions (see Appendix Fig. A1).

As shown in Fig. 2, the results indicate that *γ* increases with SPIB iteration for all systems converging toward 1, indicating the improvement during the SPIB cycles. Despite the overall upward trend in the plots, we note the final SPIB iteration in Fig. 2 does not always converge to the highest *γ*. In subplot (a) Abl kinase WT–Imatinib, for instance, we extended several iterations beyond the point where optimal RC was chosen from previous work of Ref. 36 to see if *γ* would continue to rise toward 1. However, as the results demonstrate, simply prolonging SPIB iterations does not guarantee the highest *γ*. This indicates that indefinite retraining does not necessarily yield an optimal RC in several systems. This is however, not a problem in practical terms as one can work with the RC with the highest *γ* value from all iterations.

**FIG. 2:**
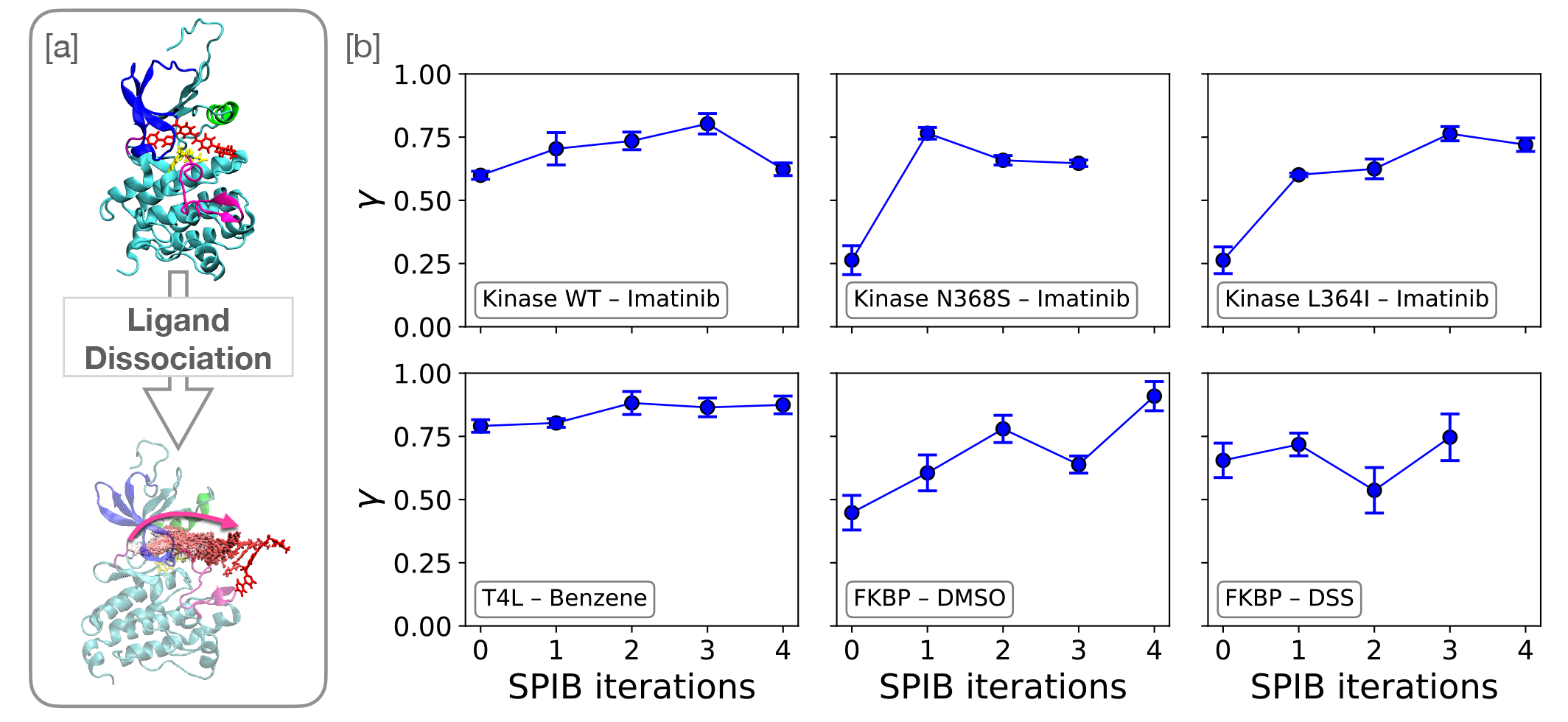
[a] Schematic of Imatinib dissociation from the Abl-kinase. [b] Values of the EATR metric *γ* over SPIB iterations for six protein–ligand systems while fixing *k*0 to the experimental value. ABL kinase Wild Type (WT) - Imatinib, ABL kinase mutant N368S - Imatinib, ABL kinase mutant L364I - Imatinib, T4 Lysozyme - Benzene, FKBP - DMSO, FKBP - DSS. Error bars denote standard deviations of fitting uncertainty in *γ*.

### B. *γ* correlates with 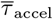

Figure 2 shows an increase in the EATR metric *γ* over successive SPIB iterations. In contrast, our earlier study demonstrated that the mean accelerated time denoted as 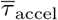 (average of the accelerated time, computed directly, without fitting to a Poisson distribution, to facilitate straightforward comparison) decreases as the RC is refined. Taken together, these opposing trends point to an inverse relationship between *γ* and 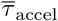, as illustrated for all six protein–ligand systems in Fig. 3 (color scale denotes SPIB iteration index). To replace this empirical observation with a more rigorous argument, here we derive an analytical expression that links *γ* to the mean accelerated time 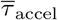 and then compared the predictive quantities against the fitted *γ* values. Starting from the EATR formalism and isolating the terms involving *γ*, we obtain

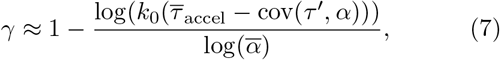

which can be further reduced to

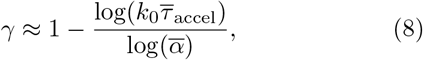

when the covariance term cov(*τ* ^′^, *α*) is negligibly small compared to 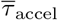. The detailed derivation can be found in the Appendix Sec.IV.

**FIG. 3:**
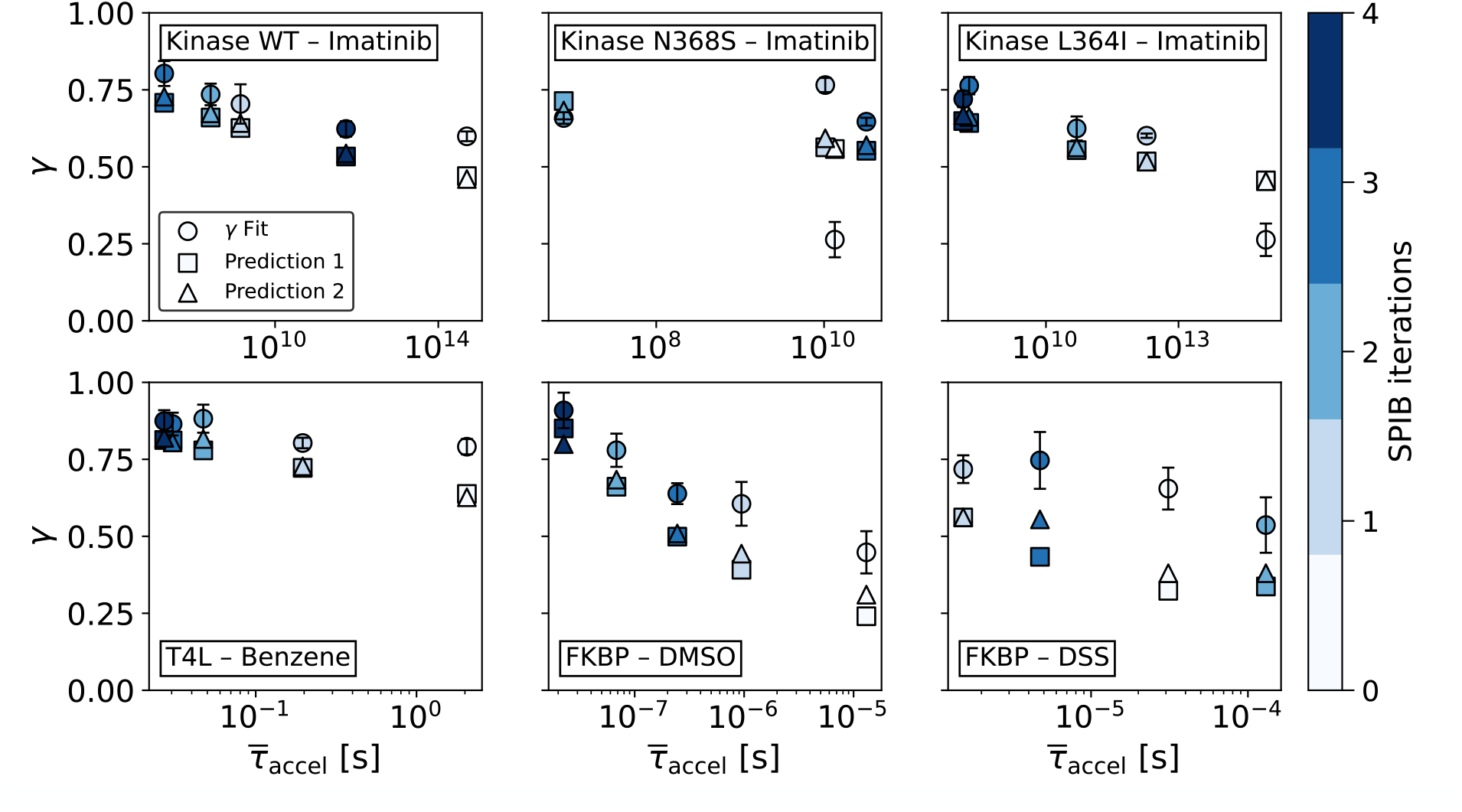
Inverse correlation between *γ* average accelerated time 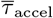 across SPIB iterations. Scatter plots are shown for six benchmark systems (arranged from top left to bottom right): Kinase WT–Imatinib, Kinase N368S–Imatinib, Kinase L364I–Imatinib, T4 lysozyme–Benzene, FKBP–DMSO, and FKBP–DSS. Circles denote *γ* values obtained from the original EATR fit, squares (“prediction 1”) show *γ* evaluated with Eq. 7, and triangles (“prediction 2”) show *γ* evaluated with Eq. 8. Different colors indicate the SPIB iteration, ranging from light blue (early iterations) to dark blue (later iterations). Error bars denote standard deviations of fitting uncertainty in *γ*.

To assess how well the analytical expressions reproduce the fitted results, Fig. 3 compares the *γ* values predicted by Eq. 7 (squares, “prediction 1”) and Eq. 8 (triangles, “prediction 2”) with the values obtained from direct EATR fitting approach (circles). The two analytical approximations practically overlap, confirming that cov(*τ* ^′^, *α*) is indeed small in most cases. Although both predictions show some, system-specific deviations from the fitted *γ*, they capture the same overall trend, demonstrating that either expression can serve as a fast diagnostic tool when a full survival-curve fit is impractical.

Collectively, the results demonstrate the reciprocal relationship between *γ* and 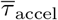. In early iterations (*γ* ≈ 0.25 − 0.5) the RC is non-optimal, whereas in later iterations (*γ* ≈ 0.7−0.9) the RC captures much better the true slow mode and the required acceleration is minimal. The consistency of this reciprocal relationship across all six systems confirms that iterative SPIB systematically improves RC quality and kinetic accuracy, and demonstrates that *γ* is, in fact, a robust metric for evaluating RC performance.

### C. Effect of bias deposition pace on the RC

As an extension to the previous analysis, we examined whether the EATR metric *γ* depends on the frequency with which Gaussian hills are deposited when the RC is held fixed. This is of immense practical utility as being able to add bias more frequently while still being able to recover accurate kinetics could have a non-linear effect on computational efficiency. We selected the optimal RC for each system, i.e., the one associated with the minimum average accelerated time 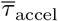 over all SPIB iterations, and carried out short infrequent-metadynamics simulations across a range of bias deposition paces.

Fig. 4 presents *γ* as a function of deposition pace for three representative systems: T4 Lysozyme–benzene, FKBP–DSS, and FKBP–DMSO. The prefactor *k*_0_ was kept fixed so that the effect of bias frequency on computed *γ* could be isolated. As 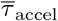 approaches the experimental residence time (gray dashed line), *γ* converges toward 1. Increasing the deposition pace up to roughly 5000 steps (i.e., decreasing the frequency of bias deposition) tends to produce a monotonic rise in *γ*; beyond this threshold, *γ* no longer improves across all three systems.

**FIG. 4:**
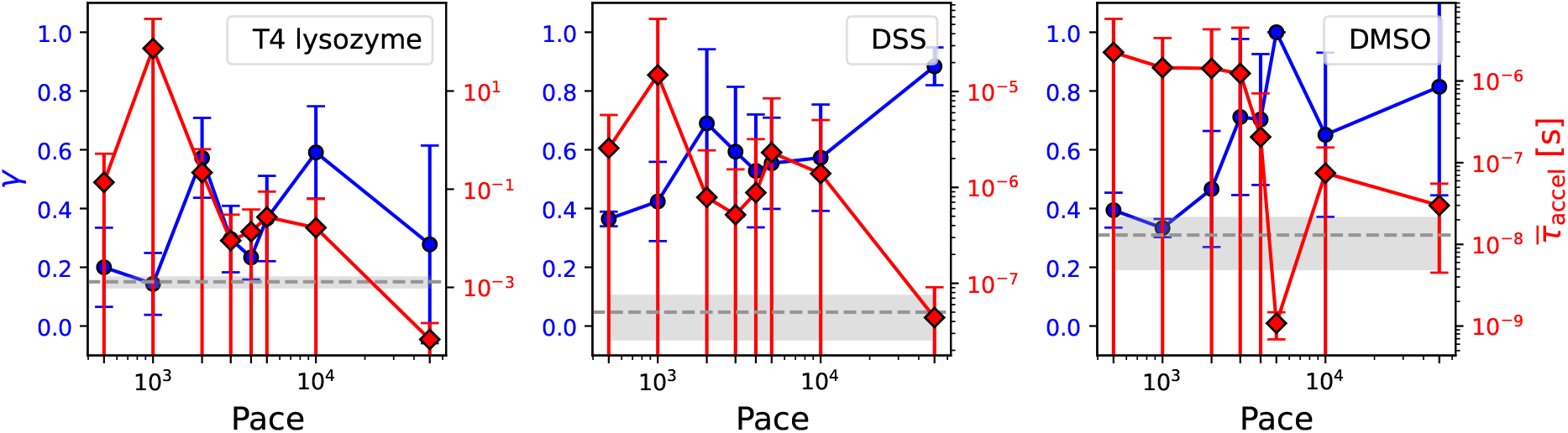
EATR *γ* metric (blue circle, left axis) and mean accelerated time ⟨*τ*⟩ (red diamond, right axis, log scale) as functions of bias-deposition pace for three protein–ligand systems. (a) T4 Lysozyme–Benzene (b) FKBP–DSS (c) FKBP–DMSO. The gray dashed line represents the experimental residence time, and the shaded gray band indicates the error margin. Error bars represent standard deviations of the *γ* and the 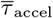.

From these findings, two conclusions follow: (i) very frequent bias deposition (pace ≤1000=2 ps) drives *γ* toward 0, leading to a suboptimal RC for kinetic studies (ii) Within a practical pace range (≈ 5000–50000), *γ* remains largely invariant to the deposition frequency, underscoring its robustness as a metric for assessing RC quality. These results are consistent with the trends observed in Ref. 38.

## IV. CONCLUSION

In this study, we performed a systematic evaluation of multiple iterations of an approximate RC derived from the SPIB framework for obtaining unbiased kinetics using iMetaD using the *γ* metric of EATR. Two questions motivated the work: (i) does iterative SPIB training measurably improve RCs, and (ii) can the *γ* metric from the EATR approach provide a good score for the evaluation? Across six different protein–ligand benchmarks, we observed an increase in *γ* along with a corresponding decrease in the mean accelerated time 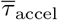 in each SPIB iteration. Importantly, we found that the quality of an RC as measured by *γ* was not strongly dependent on the pace of bias deposition, demonstrating robustness. Our reported approximate formula for CV quality (Eq. 8) is a good heuristic for whether the chosen CVs are working well to predict kinetics, but requires some knowledge of *k*_0_. Moreover, if successive iterations of an RC do not show an inverse relationship between *γ* and 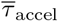, then the RC should be treated with caution. Examining these two complementary metrics side by side thus provides a straightforward, orthogonal strategy for evaluating RC quality.

We identified an important limitation. As discussed in Sec. III A, the value of *γ* is sensitive to the choice of the reference rate *k*_0_, so the parameter should be selected with care. Nevertheless, even when *k*_0_ or the bias-deposition frequency is varied, *γ* preserves the relative ordering of the RC quality. RCs that capture the slow modes better yield *γ* values closer to 1.

In general, it should be possible to directly apply EATR to iMetaD data to obtain both a good estimate of *γ* and *k*_0_. While this works well for some cases, in other cases, a low value of *γ* is obtained by this approach even though the simpler 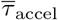 is relatively close to the experimental value. This may simply reflect a limitation of EATR in the case where limited data is available for prediction (i.e. tens of simulations to fit a rate). We are now developing an alternative approach based on the EATR formalism for obtaining even more reliable estimates of rate constants and *γ* through a combination of data from OPES-MetaD applied with multiple biasing strengths^12^, which may alleviate this problem, but that requires further investigation.

Overall, our framework is broadly applicable. First, beyond its use as a scoring metric, *γ* can be monitored during SPIB training to adaptively optimize the RC and to serve as a stopping criterion, to avoid unnecessary computation. Second, when a full fit of the *γ* metric is impractical, the predictive equations we derived offer a rapid diagnostic for assessing relative RC quality. Finally, the approach extends beyond protein–ligand kinetics to systems undergoing folding–unfolding transitions and even intrinsically disordered proteins, where suitable reference coordinates are elusive.

## Acknowledgments

This work was supported by NIH/NIGMS under award number R35GM142719 (P.T. and S.L.) and under award number R35GM138312 (G.M.H. and N.M). We thank UMD HPC’s Zaratan and NSF ACCESS (project CHE180027P) for computational resources. This work was supported in part through the NYU IT High Performance Computing resources, services, and staff expertise, and calculations were partially executed on resources supported by the Simons Center for Computational Physical Chemistry at NYU (Simons Foundation Grant No. 839534). P.T. is an investigator at the University of Maryland-Institute for Health Computing, which is supported by funding from Montgomery County, Maryland and The University of Maryland Strategic Partnership: MPowering the State, a formal collaboration between the University of Maryland, College Park, and the University of Maryland, Baltimore. The Flatiron Institute is a division of the Simons Foundation. G.M.H. and N.M. thank the Center for Computational Mathematics at the Flatiron Institute for their hospitality while a portion of this research was carried out.

## Code availability statement

A Python implementation for different *γ* evaluation methods is available on GitHub repository: https://github.com/hocky-research-group/SPIB_EATR_paper_data.

## APPENDIX

### Unbiased rates used in calculating *γ*

**TABLE T1:** The fixed values of the experimental residence times used in EATR to determine *γ* during the SPIB iterations.

| System | Transition Time (s) |
| --- | --- |
| FKBP-DMSO | $13 \times 10^{-9}$ |
| FKBP-DSS | $50 \times 10^{-9}$ |
| T4 Lysozyme L99A-Benzene | $1.3 \times 10^{-3}$ |
| Abl Kinase WT-Imatinib | 1200 |
| Abl Kinase N368S-Imatinib | 400 |
| Abl Kinase L364I-Imatinib | 700 |

### Fitting *k*_0_ in EATR

We can perform a global fit for the unbiased rate *k*_0_ in addition to *γ* by calculating the sum of square errors for each (*k*_0_, *γ*) pair, then taking the minimum. This gives the results in Fig. A1.

### Derivation of a relation between *γ* and 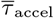

Considering the observed relationship between the average accelerated time and *γ*, we provide an argument for why this relationship should exist. We first connect the average accelerated time 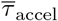 to the average biased simulation time 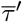:

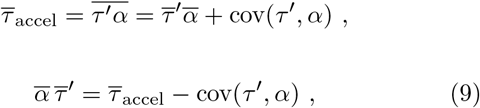

where cov(*X, Y*) is the covariance between *X* and *Y*.

We now wish to demonstrate a connection between 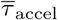 and *γ*. Using the relationship between the biased rate and the biased simulation time, 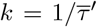, and applying Eq. 6, we can rewrite the left hand side of Eq. 9:

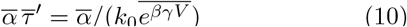

To solve for *γ*, we note that we can approximately take *γ* outside of the ensemble average for large values of *e*^*βV*^ (when 0 *< γ <* 1):

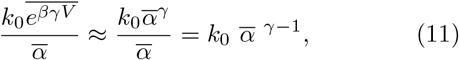

because 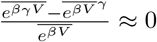.

Combining Eqs. 10 and 11 we get,

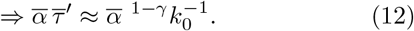

Equating Eqs. 9 and 12 gives us

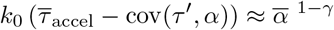

and taking the logarithm on both sides and solving for *γ*, we obtain the following two equations presented in Sec. III B:

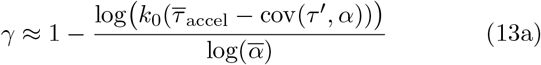

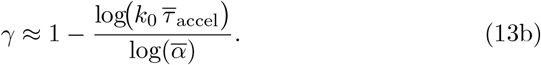

**FIG. A1:**
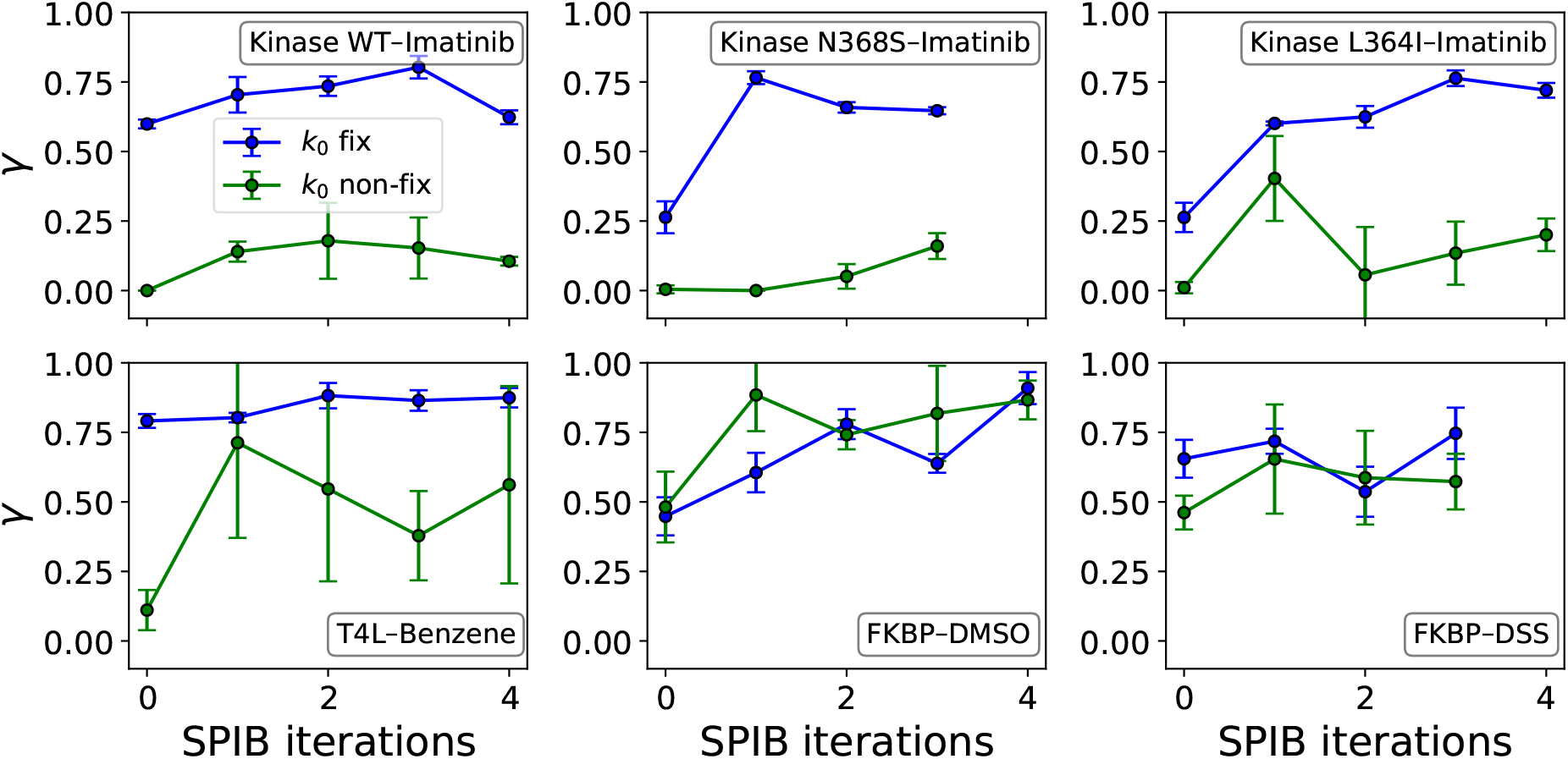
Comparison of fitted *γ* values with fixed *k*_0_ versus free *k*_0_. Each subplot shows *γ* plotted against SPIB iteration for six different protein–ligand systems. Error bars denote the standard deviation of the fitting uncertainty in *γ*. Blue plot represents *γ* value in which *k*_0_ was held fixed at its experimental value (as in Fig. 2), while green plot represents *γ* value in which both *k*_0_ and *γ* were treated as free parameters.

**FIG. A2:**
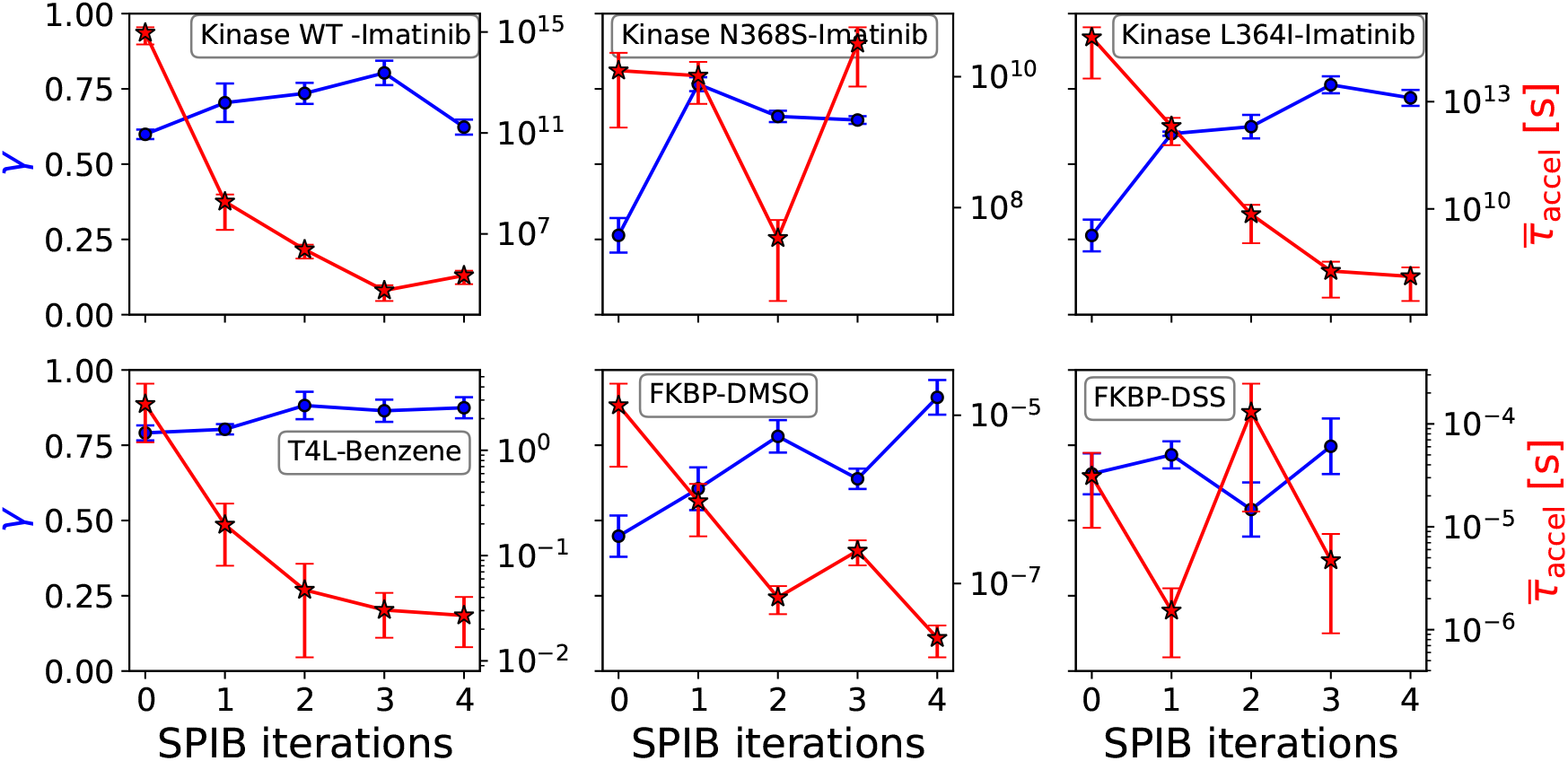
Same data as Fig. 3 represented in a different way. Here, results of the EATR metric *γ* (blue circle, left axis) and the mean accelerated time 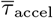 (red star, right axis, log scale) over SPIB iterations for six protein–ligand systems with *k*_0_ fixed to the experimental value. Top left to bottom right: ABL kinase Wild Type (WT) - Imatinib, ABL kinase mutant N368S - Imatinib, ABL kinase mutant L364I - Imatinib, T4 Lysozyme - Benzene, FKBP - DMSO, FKBP - DSS. Error bars denote standard deviations of fitting uncertainty in *γ* and standard deviations in 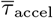.

